# Live triatomine bug, vector of *Trypanosoma cruzi* (Chagas disease), found engorged in Lisbon hotel room: A first for Portugal and for Europe

**DOI:** 10.64898/2026.02.26.708074

**Authors:** Jennifer K. Peterson, Alexander Kelley, Trinity Antoszewski, Madeline Brown, Hanna Cortes, Peyton I. Easton, Grace Ferry, Thoburn Freeman, Cate Freiwald, Erik Hagen, Hunt Kinnaird, Luna Lewin, Malaki Lewis, Jessie Mcnulty, Natalie Moore, Erin Mullis, Stella Pettit, Luke Schultz, Sarah Sharp, William Stocker, Jillian Tunstall, Jader de Oliveira

## Abstract

Triatomine bugs are blood feeding insects that transmit the parasite *Trypanosoma cruzi*, causative agent of Chagas disease. The bugs are found primarily in the Americas with a few species in Asia and Africa. Here we report the first case of a live triatomine bug in Europe, found in a Lisbon hotel room. In August, 2025, the hotel room occupants discovered a triatomine bug perched on the headboard of their bed. Upon capture, bright red blood emerged from the bug; the occupants suspected that it had bitten them during the night. The bug was identified morphologically as triatomine species *Hospesneotomae protracta*, which was confirmed molecularly. *Hospesneotomae protracta* is native to the southwestern United States where it is a competent *T. cruzi* vector. *Trypanosoma cruzi* was not detected in this specimen. Although this case likely represents an accidental importation, it illustrates the ease with which disease vectors can be unknowingly transported globally. Ergo it is crucial to document and share these findings to prevent introductions of non-native arthropods of medical importance.

## Introduction

Triatomine bugs (Hemiptera: Reduviidae: Triatominae) are obligate hematophagous insects found primarily in the New World.^1^ Commonly known as “kissing bugs,” the insects are of public health significance due to their ability to harbor and transmit the causative agent of Chagas disease, *Trypanosoma cruzi*. Chagas disease is a chronic illness that, when left untreated, can lead to serious cardiac and/or gastrointestinal alterations.^2^ An estimated 6–7 million people worldwide are infected with *T. cruzi*.^3^

Triatomine bug diversity and abundance are concentrated in tropical and subtropical regions of the Americas (referred to as ‘New World’ species).^4^ A handful of triatomine bug species are found in in Africa, Asia, Australia and the South Pacific (‘Old World’ species), but not in continental Europe.^5^ Notably, Old World Triatominae are not believed to harbor or transmit *T. cruzi*. The sole report of a triatomine bug in Europe occurred in 2022 when a single dead specimen of Old World species *Triatoma rubrofasciata* was found in Spain in a commercial shipment from China. Here, we present the first case of a live, free-living New World triatomine bug species in Europe, which was found engorged with fresh blood in a luxury hotel room in Lisbon, Portugal. The specimen was likely imported accidentally and does not represent an introduced European triatomine bug population. Nonetheless, the case is illustrative of the ease with which disease vectors can be unknowingly transported globally.

### Case description

On the morning of August 26^th^, 2025, a live triatomine bug (Fig. 1) was found in a 10^th^ floor luxury hotel room in Lisbon, Portugal. The bug was found alive on the headboard of the hotel bed and upon capturing the insect with a tissue, bright red blood emerged from below its pronotum, suggesting that it had recently fed (Fig. 1). The bug was discovered by the hotel room occupants, a couple visiting Lisbon from the east coast of the United States (US). They were in Lisbon as part of a pre-trip extension prior to taking a river cruise. They had arrived the day before (August 25^th^) and spent one night in the room. They believed that one of them had been bitten by the bug during the night because one of them had woken up with a dark brown streak on their arm and finger, which they believed to be the insect’s feces.

**Figure 1.**
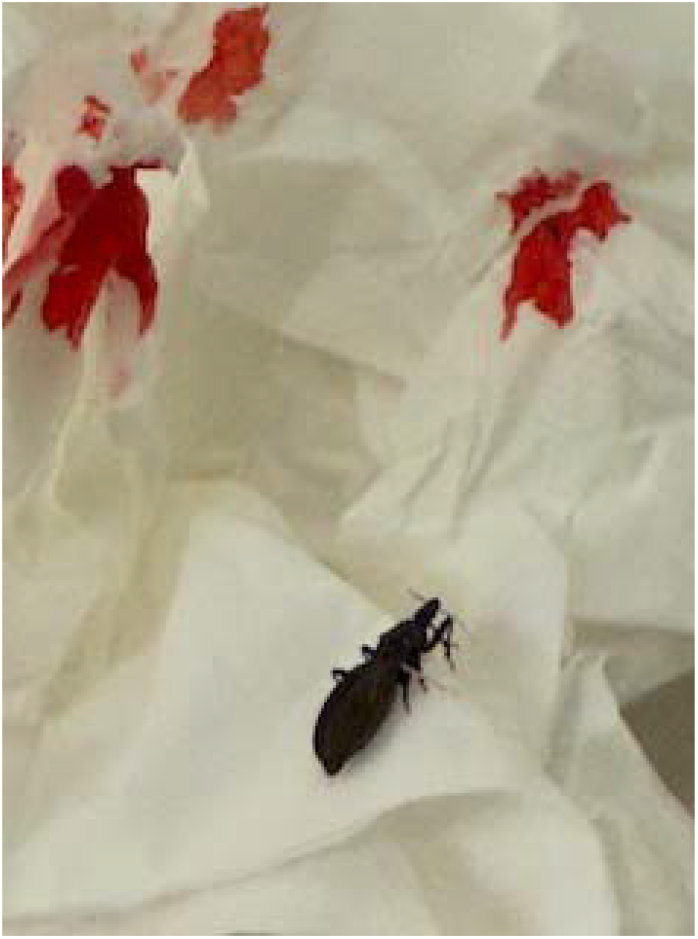
Triatomine specimen moments after capture, resting on the tissue used to absorb the blood that emerged from its abdomen. Alt text: Triatomine bug specimen resting on tissue with bright red blood on it.

The couple reported the incident to the hotel desk receptionist and hotel manager, showing them photographs of the bug (Fig. 1). The hotel staff was apologetic and surprised, stating that they could not imagine how the bug had gotten there. The couple was moved to a different room and compensated for their troubles. They stayed in the hotel (in a different room) for three additional nights and did not encounter any more triatomine bugs.

The hotel was in high-rise building in an upscale area of Lisbon. The room occupants who found the bug reported that they observed several other cruise guests staying in the same hotel, suggesting that it is an establishment that caters to international tourists. They reported the condition of the hotel as ‘nice but needing cleaning.’ The hotel is advertised on its website as a five-star establishment.

Upon returning to the US, the couple reached out to their primary care physician because they were concerned about possible *T. cruzi* exposure and Chagas disease. The physician dismissed their concerns, after which they requested assistance from the Infectious Disease Entomology Across Scales (IDEAS) lab in the Department of Entomology and Wildlife Ecology at the University of Delaware. They sent the specimen to the IDEAS lab, where it was identified to species and tested for *T. cruzi* infection. They also consulted an infectious disease specialist, who ordered a script for the couple was to be screened *T. cruzi* antibodies, which they did at a commercial diagnostic laboratory (Labcorp). Available information for Labcorp indicates that they use Indirect Fluorescent Antibody (IFA) testing the Alinity s Chagas test^6^, which is a chemiluminescent microparticle immunoassay (CMIA) manufactured by Abbott™. Antibodies were not detected in either spouse.

Despite having undergone two transatlantic journeys, the specimen was relatively intact, with just its antennae and parts of the legs missing or loose (Fig 2; all legs fell off during positioning for lateral view photo). The specimen was identified as an adult female *Hospesneotomae protracta* Uhler, 1894 (formerly *Triatoma protracta*^*7*^). The species is native to the southwestern US and Northwest Mexico, and an important North American vector of *T. cruzi*. Species identification was made using a consistent set of external morphological characters, following the classical keys by Lent & Wygodzinsky^8^ and the updated taxonomic framework proposed by Paiva et al.^7^ In brief, the identification was based on body size (males 13–19.5 mm, females 15–23 mm), a trapezoidal pronotum that is smooth and laterally carinate with a slightly elevated anterior lobe lacking tubercles, and a rugose posterior lobe with rounded humeral angles. The head is about twice as long as wide, distinctly convex dorsally, and shows a clear arcuate transverse impression at the base of the clypeus, visible in both dorsal and lateral views. The anteocular region is much longer than the postocular region, the genae taper apically without reaching the clypeus, and the eyes are relatively small, reaching the level of the upper or lower surface of the head in lateral view. Antenniferous tubercles are positioned near the middle of the anteocular region.

**Figure 2.**
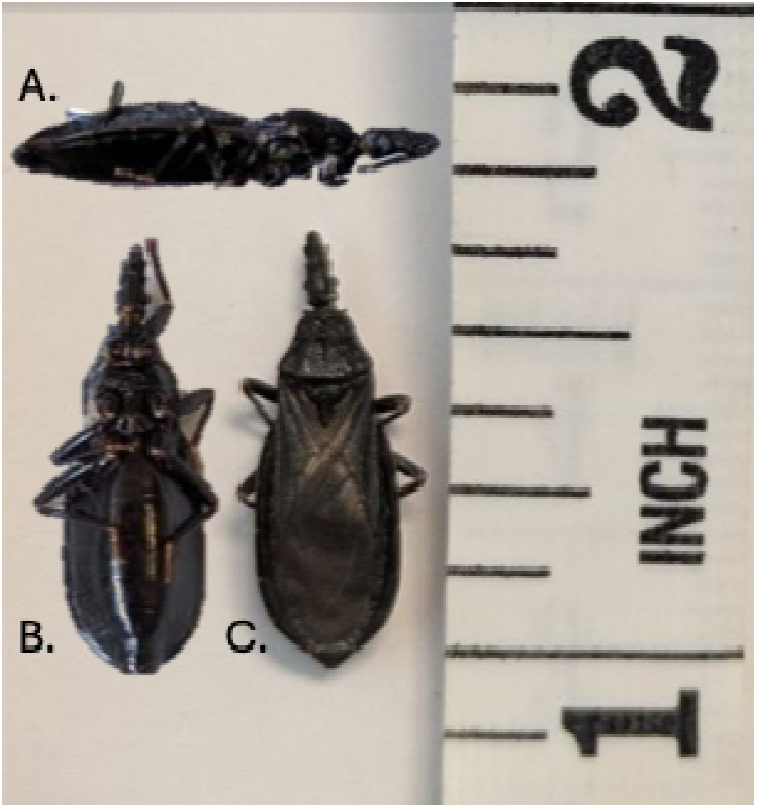
Lateral (A), ventral (B), and dorsal (C) views of *H. protracta* specimen found in Lisbon hotel room. Alt text: Three different views (top, side, and bottom) of triatomine specimen found in Lisbon hotel room.

**Figure 3.**
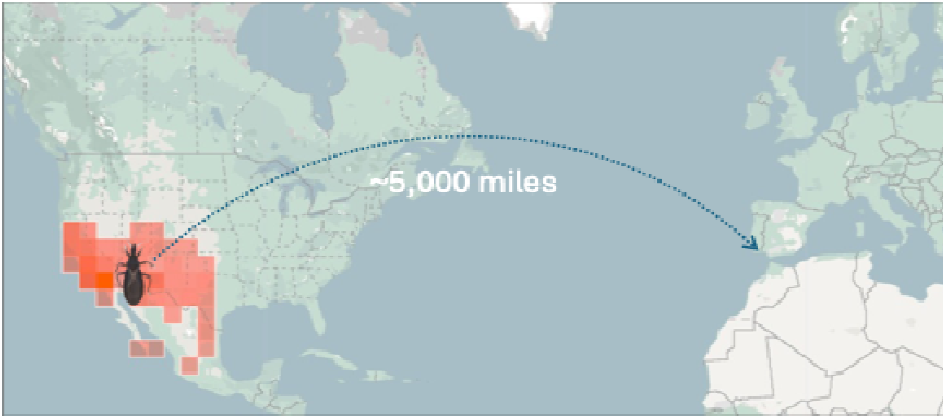
Possible route traveled by specimen. Approximate geographic range of *H. protracta* shown in orange. Data and map from inaturalist.com, which integrates with Google maps. Alt text: Flat map of the northern hemisphere showing all of the United States and Western Europe. The southwestern corner of the United States is highlighted in orange to indicate the native range of H. protracta. A curved, dotted line from the center of the southwestern US range (approximately New Mexico) to Lisbon, Portugal indicates the potential path traveled by the specimen.

Molecular confirmation of the species was carried out through PCR and sequencing, as described previously.^9–11^ DNA extracted from the insect’s lower abdomen was amplified in a conventional PCR using the cytb-intF/R primers (5’-YCT ACT ATC CGC GGT TCC TTA-3’ and 5’-ATA CTA TTG CAA TTA CTC CTC CTA-3’, respectively), which amplify a 443 base pair fragment of the triatomine mitochondrial cytochrome *b* (cyt B) gene. Amplicons were Sanger sequenced and sequences were aligned in Genbank using the BLAST function.^12^ The top eight alignments matched the species *Hospesneotomae protracta* with the top hit having 96.53% identity, 98.00% query cover, and an e-value of 0.00.

*Trypanosoma cruzi* DNA was not detected in the specimen, which was screened for the parasite in a duplex qPCR run in triplicate.^13^ Briefly, DNA extracted from the specimen was amplified using the cruzi1 and cruzi2 primers (5’-ASTCGGCTGATCGTTTTCGA-3’ and 5’-AATTCCTCCAAGCAGCGGATA-3,’ respectively) and the cruzi 3 probe (5’-FAM-TTGGTGTCCAGTGTGTG-NFQ-MGB-3’), which target a 165 bp sequence of *T. cruzi* satellite DNA. A 163 bp sequence of triatomine 12s ribosomal RNA was targeted in the same reaction using the P2B (5’-AAAGAATTTGGCGGTAATTTAGTCT-3’) and P6R (5’-GCTGCACCTTGACCTGACATT-3’) primers and the Triat probe (5’-VIC-TCAGAGGAATCTGCCCTGTA-NFQ-MGB-3’) as an internal amplification control (IAC). The IAC successfully amplified triatomine DNA, with a strong positive signal and critical threshold values of 17 for all three replicates, providing further taxonomic confirmation that the specimen is a triatomine bug.

All details and data for the specimen were shared with the Center for Vectors and Infectious Diseases Research (CEVDI) at the Portuguese National Institute of Health Doctor Ricardo Jorge (INSA).

## Discussion

Here we present the first case of a live, New World triatomine bug species in Europe. We detected and identified *Hospesneotomae protracta*, a species from North America, in a hotel in Lisbon, Portugal. This individual was likely an accidental importation that was transported overseas passively, which is common for smaller disease vectors such as mosquitoes,^14^ but just a handful of modern day reports exist of accidental overseas triatomine bug transport. In pre-Columbian populations, highly domesticated triatomine species such as *Triatoma infestans* and *Rhodnius prolixus* are thought to have been passively dispersed during human activities and overland movement of their goods and animals^15– 18^ and *T. rubrofasciata* is hypothesized to have dispersed with ship rats on ancient maritime trade routes.^19,20^ However, modern reports of passive overseas dispersal are rare. In the US, two *T. ryckmani* individuals were intercepted at the Miami airport in 1970 on a Bromeliad sent from Guatemala.^21^ More recently, as mentioned above, a dead *T. rubrofasciata* specimen was discovered in a box of commercial goods in Spain in 2022.^22^ The specimen had hitchhiked to Spain from China, where the triatomine bugs are not believed to harbor *T. cruzi*. A North American confamiliar (in the Reduviidae family) species, *Zelus renardii*, was first recorded in Greece in 2011 and believed to have been introduced through imported plants. *Zelus renardii* preys on beneficial and harmful crop insects and is not a human disease vector. The species was found in Portugal for the first time in 2020, expanding by an estimated 40km per year,^23–25^ illustrating the speed with which introduced insects can spread once the population is seeded. Interestingly, *H. protracta* is one of just a handful of Triatominae species that glues their eggs onto the substrate upon which they oviposit, which lends itself to passive dispersal.^26^

We cannot ascertain the exact geographical origin of the *H. protracta* specimen or the way in which it was transported to Lisbon, but it probably came from somewhere in the southwestern United States (US) and was transported in someone’s personal belongings or a shipment of commercial goods.

Portugal is an increasingly popular destination for American tourists and American exports; in 2024, 2.3 million US tourists visited Portugal, up 12% from the previous year, while 244 million dollars of agricultural goods were exported to Portugal from the US in 2022.^27,28^ The states of California and Texas, both endemic for *H. protracta*, are among the top six US states with the highest exports to Portugal.

Behavioral characteristics specific to *H. protracta* may have contributed to the realization of the specimen’s transatlantic journey. First, the species is native to arid and semi-arid regions of the southwestern United States and northwestern Mexico, adapted to desert climates with hot summers and cool, dry winters. A 20-year study of *H. protracta* populations in Griffith Park, Los Angeles CA found that peak movement occurred during the months of July and August, aligning with the time of year (August) in which the bug was found in Lisbon. These dispersals occurred when temperatures were between 60-76°F,^29^ which is close to the average temperature of the cargo hold of a commercial jet, which is pressurized and kept around 65°F.^30^ If the specimen was transported to Portugal in someone’s personal belongings, this adaption to cool climates may have helped it to survive the nine hour (or more) flight to Lisbon from the southwestern US. Additionally, the species is closely associated with the grassy, inner breeding cavities of woodrat middens, thus accustomed to tight spaces.^31^ Indeed, many Triatominae species, including *H. protracta* exhibit a ‘sit and wait’ nest predation strategy where they remain in tiny spaces for long periods of time awaiting their next blood meal.^31^ These species have a high starvation capacity (weeks to months), which would also contribute to their survival on long-haul trips and the time spent afterward awaiting a blood meal.^32,33^ When given the opportunity to feed however, *H. protracta* are known to be aggressive feeders, taking advantage of available bloodmeals in laboratory settings within 30 to 60 seconds.^34^ In the wild, they will enter human homes and feed opportunistically on humans and their animals, as the specimen described here did, and several reports of *H. protracta* bites exist in its native geograhic range.^29,34–36^

Although *T. cruzi*, was not detected in the *H. protracta* specimen reported here, the species is a *T. cruzi* competent species. In a study of 1,759 individuals collected in Los Angeles, CA between 1941-1967, Wood et al. observed trypanosomes in the feces of 593 individuals, revealing a prevalence of 33.1%. A PCR-based analysis likely would have revealed an even higher prevalence, as parasites are not always visible in triatomine feces. A more recent study of *T. cruzi* infection in *H. protracta* in California found a prevalence ranging between 18.7% (26/139) and 36.4% (8/22), indicating that the parasite still actively circulates in the region.^35^

Despite being competent for the *T. cruzi* infection, the epidemiological risk of *H. protracta* transmitting *T. cruzi* to humans is considered low due the limited interaction of *H. protracta* with humans.^35^ *Trypanosoma cruzi* is transmitted in the vector’s excrement, (not its saliva), meaning that the parasite is *not* injected into the host during the blood meal. One study found that an average of five percent of lab-reared adult *H. protracta* defecated during feeding and an additional seven percent defecated within one minute of repletion, suggesting that the species will defecate on its host.^36^

Finally, there is also a very low probability of an imported uninfected triatomine bug acquiring the infection in the non-endemic region. There are an estimated 125,000 people infected with *T. cruzi* in Western Europe.^37^ In Portugal, the *T. cruzi* infection prevalence among migrants from Chagas-endemic regions is unknown; the World Health Organization estimate is between 900 and 89,900^38^ with the Chagas disease underdiagnosis rate estimated to be over 99%.^39^ In addition, climate modeling suggests that several areas of Portugal and Southern Europe are climatically suitable for some triatomine species, should a population be introduced.^40^ These data serve to highlight the importance of surveillance for triatomine importations in non-endemic regions.

## Conclusion

The case presented here highlights the ease with which portable and sturdy Triatominae species can be passively transported long distances to non-endemic regions. Considering the incidence in the context of ever-increasing global trade and tourism combined with climate change and expansion of habitat suitability for these vectors, this case illustrates the importance of remaining vigilant to the entry, establishment, and spread of potentially harmful insect species.

## Acknowledgements

The authors are indebted to the hotel room occupants represented in this paper for their collaboration throughout the process described above. Access to the sequencing and informatics was supported by the Institutional Development Award (IDeA) from the National Institute of Health’s National Institute of General Medical Sciences under grant number P20GM103446. We would also like to acknowledge support from USDA Hatch (DEL00854 and NE2443). The author (JO) acknowledges the financial support provided by CAPES (Finance Code 001).

